# Exploring the quality of protein structural models from a Bayesian perspective

**DOI:** 10.1101/2020.07.27.223818

**Authors:** Agustina Arroyuelo, Jorge A. Vila, Osvaldo A. Martin

## Abstract

We explore how ideas and practices common in Bayesian modeling can be applied to help assess the quality of 3D protein structural models. As the word *model* is used in both Bayesian Statistics and Protein Science, throughout this article we deliberately use the word *model* to discuss statistical models and *structure* to discuss protein 3D models, thus avoiding potential confusions. The basic premise of our approach, is that the evaluation of a Bayesian statistical model’s fit may reveal aspects of the quality of a structure, when the fitted data are related to protein structural properties. Therefore, we fit a Bayesian hierarchical linear model to experimental and theoretical ^13^C^*α*^ Chemical Shifts. Then, we propose two complementary approaches for the evaluation of such fitting: 1) in terms of the *expected differences* between experimental and posterior predicted values; 2) in terms of the *leave-one-out cross validation point-wise predictive accuracy*. Finally, we present visualizations that can help interpret these evaluations. The analyses presented in this article are aimed to aid in detecting problematic residues in protein structures. The code developed for this work is available on: https://github.com/BIOS-IMASL/Hierarchical-Bayes-NMR-Validation.

## Introduction

Bayesian statistics offers very suitable theoretical advantages for developing models involving bio-molecular structural data and it has been applied in numerous tools and methods in this context [1–6]. Furthermore, Bayesian methods are capable of accounting for errors and noise of variable source and nature, which is suitable for working with bio-molecular experimental data [7].

In statistics, partially pooling data means to separate observations into groups, while allowing the groups to remain somehow linked in order to *influence each other*. In a Bayesian setting, such sharing is achieved naturally through hierarchical modeling. In hierarchical models (also called multilevel models), parameters of the *prior* distributions are shared among groups, inducing dependencies and allowing them to effectively *share information* [8–10]. Advantages of hierarchical Bayesian modelling include obtaining model parameter estimates for each group as well as for the total population. Also, using common *prior* distributions helps preventing the models over-fit [11].

In this study, we fit Bayesian models to experimental ^13^C^*α*^ Chemical Shifts.We focus specifically on ^13^C^*α*^ Chemical Shifts because they have proven to be informative on protein structure at the residue level and have been used in determination and validation in previous work by us and others [12–15]. Another promising aspect of working with ^13^C^*α*^ Chemical Shifts is that their theoretical counterparts, can be computed with high accuracy using quantum-chemical methods [13].

Bayesian model comparison and evaluation is standard in Bayesian applications as it constitutes a very important part of the Bayesian workflow. A variety of methods have been proposed for this task, helping Bayesian practitioners evaluate, critique and ultimately understand their models [16]. In the present work, we propose two complementary approaches for the evaluation of protein structures. Both are related to different ways of analysing the results of a Bayesian hierarchical linear model that links experimental and theoretical ^13^C^*α*^ Chemical Shifts. For the first approach, we compare structures in terms of their residuals (i.e. the difference between the observed and posterior predicted values). To ease the comparison, we put the residuals in the context of reference densities which are pre-computed from a data set of high quality protein structures (see Methods and Software section for details). In the second approach, we evaluate the statistical model’s fit in terms of its out-of-sample predictive accuracy, i.e. the predicted accuracy computed from data not used to train the model. The out-of-sample predictive accuracy can be estimated using leave-one-out cross-validation, which requires to re-fit a model *n* times, with *n* being the size of the data set (i.e. the number of ^13^C^*α*^ Chemical Shifts). As this can be too costly and cumbersome, in this work we use an alternative; the Pareto-smooth-importance sampling leave-one-out cross validation (LOO for short) [17, 18]. LOO offers an accurate, reliable and fast estimate of the out-of-sample prediction accuracy from a single model fit. Additionally, the predictive accuracy is computed per observation, as we use ^13^C^*α*^ Chemical Shifts this is equivalent to compute the predictive accuracy per residue. This allows us to make statements of the quality of the structure at both global and per residue level.

We expect the methods and visualizations presented here to introduce Bayesian model checking tools to protein scientists. Also, we hope these methods are adopted by protein scientists to help them evaluate the quality of a given structure. This may include the structure determination process before structure deposition at the Protein Data Bank. Or even after deposition, such as evaluating a structures quality prior to further research like, for example, performing docking or template-based modeling.

## Methods and Software

### Reference data set

A reference data set of 111 high quality protein structures was obtained from the Protein Data Bank (PDB) [19]. Each structure in this set has a resolution ≤ 2.0 Å and R-factors ≤ 0.25. The structures do not contain coordinates for DNA, RNA or glycan molecules. Additionally, every structure in our set has a corresponding entry at the Biological Magnetic Resonance Bank (BMRB) from which experimental ^13^C^*α*^ Chemical Shift data was obtained [20]. Theoretical 13C^*α*^ Chemical Shift data was computed from the Cartesian coordinates of each structure in this set, using *Che*Shift-2 [15].

### Target structures

Theoretical ^13^C^*α*^ Chemical Shift data was obtained for two structures of protein Ubiquitin under PDB ids: 1UBQ and 1D3Z [21, 22]. Code id.: 1UBQ corresponds to an X-ray crystallography determined structure of Ubiquitin, while 1D3Z corresponds to an NMR determined structure. The latter contains ten different conformations. We computed and averaged the theoretical ^13^C^*α*^ Chemical Shifts for those 10 conformations. Additionally, the experimental ^13^C^*α*^ Chemical Shift set used in this analysis is taken from BMRB entry N°: 6457, and is the same experimental data set used in the NMR determination of 1D3Z. In this study, the same experimental ^13^C^*α*^ Chemical Shift set was used for both structures, but the theoretical ^13^C^*α*^ Chemical Shift set is different for each structure, given that is was obtained from the Cartesian coordinates of the PDB entries 1UBQ and 1D3Z. This particular data set construction for the target structures, allows us to compare 1D3Z and 1UBQ based solely on the differences between the 3D coordinates of their structures.

### Hierarchical linear model

A Bayesian hierarchical linear regression model was fitted to the experimental and theoretical ^13^C^*α*^ Chemical Shifts contained in the reference data set. The data was normalized before fitting by subtracting the empirical mean and dividing by the empirical standard deviation. The data grouping criterion was amino acid side-chain, i.e., the model divides the data set into 19 groups (one for each amino acid), with Cysteine being excluded given that *Che*Shift-2 does not offer reliable calculations for this amino acid. The full model is described by expression 1 using standard statistical notation and it is also represented in Figure 1 in Kruschke’s diagrams [23].

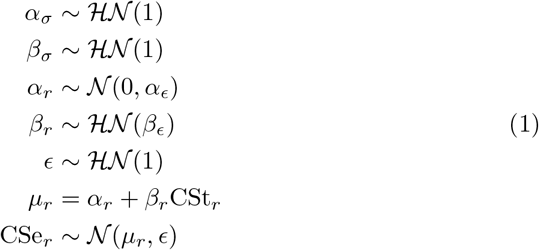

where 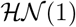 stands for half-normal distribution with standard deviation 1. 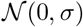 is a normal distribution with mean 0 and standard deviation *σ*. CSt_*r*_ represents theoretical ^13^C^*α*^ Chemical Shifts and CSe_*r*_ the experimental ones. The sub index *r* denotes each of the 19 groups.

**Figure 1:**
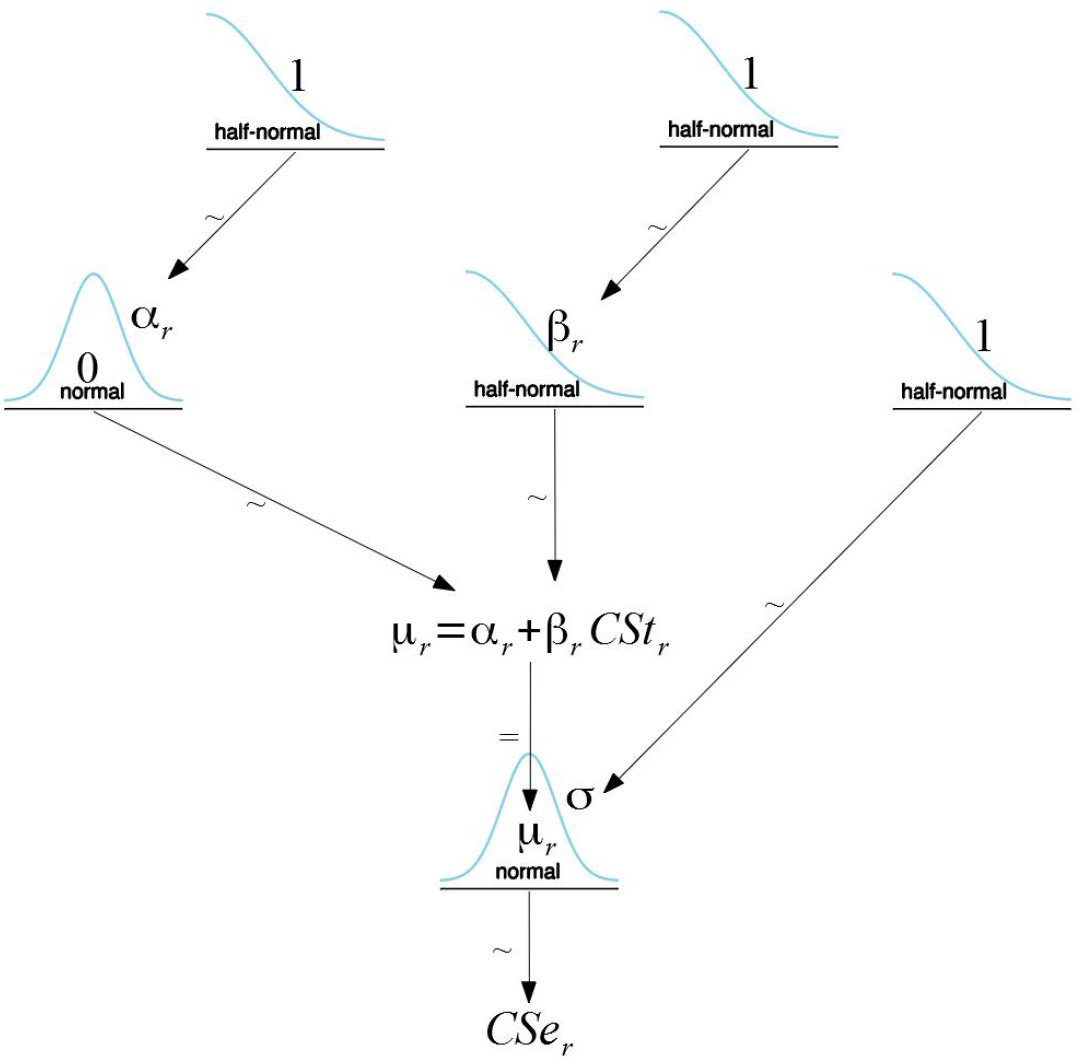
Krushke diagram representing the hierarchical linear model featured in this work.

### Computation of the posterior predictive distribution of ^13^C^*α*^ Chemical Shifts and Reference Densities

From the fitted hierarchical model, we compute the posterior predictive distribution, that is the distribution of ^13^C^*α*^ Chemical Shifts as predicted by the statistical model. We will refer to this set as corrected ^13^C^*α*^ Chemical Shifts. Then, we compute the reference densities as the difference between the corrected and experimental ^13^C^*α*^ Chemical Shifts from the reference data set for each of the 19 most common amino acids present in proteins (with Cysteine excluded as previously explained). Intuitively, the reference densities are an approximation to the expected distribution of the difference between experimental and corrected ^13^C^*α*^ Chemical Shifts. The difference between corrected and experimental ^13^C^*α*^ Chemical Shifts was also computed for the target structures, where the corrected set was defined from the posterior predictive distribution of the model fitted to the structure’s ^13^C^*α*^ Chemical Shift data.

### Model comparison and Cross Validation

When faced with more than one model for the same data it is natural to ask which model is the best at explaining the data, and more broadly, how are models different from each other and what they have in common. One way to asses a model is through its predictions. In order to do so, we can compare a model’s predictions to experimental data. If we use the same experimental data used to fit the model, i.e. we compute the within-sample error, we may become overconfident in our model. The most simple solution is to compute the out-of-sample error, this is the error that a model makes when evaluated on data not used to fit it. Unfortunately, leaving a portion of the data aside just for validation is most often than not a very expensive luxury. NMR structure validation is clearly one such example.

The log predictive density has an important role in model comparison because of its connection to the Kullback-Leibler divergence, a measure of closeness between two probability distributions [11]. For historical reasons, measures of predictive accuracy are referred to as information criteria and they are a collection of diverse methods that allow to estimate the out-of-sample error without requiring external data. In a Bayesian context one such measure is LOO as previously mentioned in the introduction [17, 18, 24]. In the next subsections, we will briefly explain some of the details related to LOO, specially those more relevant for the current work.

### LOO

The cross-validated leave-one-out predictive distribution *p*(*y_i_* | *y*_−*i*_) (or most commonly its logarithm) can be used to asses the out-of-sample prediction accuracy. In the present work this means the probability, according to the model, of observing the *i* ^13^C^*α*^ Chemical Shifts when that Chemical Shift is not included in the fitting.

Computing *p*(*y_i_* | *y*_−*i*_) can become costly as it requires to fit a model *n* times (where *n* is the data set’s size). Fortunately, the leave-one-out predictive distribution can be approximated by using importance weights. The variance of these importance weights can be large or even infinite, LOO applies a smoothing procedure that involves replacing the largest importance weights with values from an estimated Pareto distribution. For details of how this is done and why it works see Vehtari et. al. (2017) [17]. What is most important for our current discussion is that the 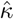 parameter of such Pareto distribution can be used to detect highly influential observations, i.e. observations that have a large effect on the predictive distribution when they are left out. In general higher values of 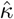 can indicate problems with the data or model, specially when 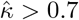 [18, 25]. In the Results section, we show how this diagnostic can be useful in the context of protein structure validation.

### LOO-PIT

PIT (Probability Integral Transform) known as the universality of the uniform distribution, states that given a random variable with an arbitrary continuous distribution, it is possible to create a uniform distribution in the interval [0, 1]. Specifically, given a continuous random variable *X* for which the cumulative distribution function is *F_X_*, then *F_X_*(*X*) ~ *Unif* (0, 1).

LOO-PIT is obtained by comparing the observed data *y* to posterior predicted data 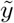. The comparison is done point-wise. We have:

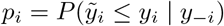

Intuitively, LOO-PIT is computing the point-wise probability that the observed data *y_i_* has a lower value than the posterior predicted data 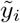. For a well calibrated model the expected distribution of *p* is the uniform distribution over the [0, 1] interval, i.e. a standard uniform distribution. Deviations from uniformity indicate different mismatches between the data and the predictions made by the model. For example a *frown shape* indicates that the predictive distributions are too broad compared with the data, while the opposite will be true for a *U-shaped* LOO-PIT density. An important advantage of using the leave-one-out predictive distribution instead of just the predictive distribution is that with the former, we are not using the data twice (once to fit the model and once to validate it) [11, 25].

### Expected Log Predictive Density

Finally, we can compare models point-wise using their Expected Log Predictive Density (ELPD), where the expectation is taken over the whole posterior. In other words, the predictions take into account the parameter’s uncertainty, as expressed by the statistical model and data. Notice that the value of the ELPD is not useful by itself, but can be used to compare the relative fit of residues within a same structure and/or to compare the relative fit of residues from two or more structures, as long as they are fitted to the same data. While the values of 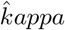 indicate how influential an observation is, i.e how much the predictive distribution changes when they are left out. The ELPD indicates how difficult is to the model to predict a particular observation.

### Software

All Bayesian models were solved with PyMC3 [26]. ArviZ was used to compute LOO, ELPD and related plots [27]. PyMOL was used to visualize 3D protein structures [28].

## Results

In this work we explore the quality of protein structures using ^13^C^*α*^ Chemical Shifts through the evaluation of a Bayesian hierarchical linear model’s fit. An important aspect of fitting this particular model, is the estimation of the effective reference value for the ^13^C^*α*^ Chemical Shifts. This is important as wrong referencing can be an issue when working with Chemical Shifts. In our study, the estimated reference value is unique for every protein in our data set. Moreover, by using a hierarchical model, we obtain a correction specific for every one of the 19 most common amino acids that constitute proteins (as already mentioned Cysteine is excluded). Said effective reference is accounted for in every posterior analysis made on the model’s fit.

We present two approaches for the Bayesian model’s fit evaluation. The first approach analyses differences between corrected and experimental ^13^C^*α*^ Chemical Shifts. This is highly appealing as the comparison is done using a familiar metric for protein scientists, specially NMR spectroscopists. The second approach, instead evaluates the model’s fit using the LOO predictive distribution, which is a general and widely accepted way to assess Bayesian statistical models [11,17,18,25]. Both methods complement each other, the first one focus on how well the corrected ^13^C^*α*^ chemical shift agrees with the expected distribution while the second approach is based on how well the model predicts the data. The combined usage of both approaches can help spectroscopists and protein scientists in general to flag problematic residues that may deserve further attention.

### Difference between experimental ^13^C^*α*^ Chemical Shifts and corrected ^13^C^*α*^ Chemical Shifts

Using reference densities on the differences between corrected and experimental ^13^C^*α*^ Chemical Shifts obtained from high quality structures can help contextualize particular differences found in any given NMR structure. Resulting in a straightforward way to asses how problematic is the resolved structure for each residue.

The reference densities for the amino acid types present in Ubiquitin are plotted in light blue in Figure 2. As expected, these distributions have zero-mean but most importantly they have different variances. Even when we did not perform any formal test to evaluate if and how these densities depart from a Gaussian distribution we can see that most of the reference densities are skewed or have more than one peak (most likely reflecting sub-populations of ^13^C^*α*^ Chemical Shift differences corresponding to *α*-helix and *β*-strands). These distributions also reflect variations among amino acids related to natural abundance as well as chemical features. For example Glycine, which is the most abundant amino acid and the one spanning broader regions on the Ramachandran plot, has the smoothest curve. Given all these particularities, comparing differences between observed and corrected ^13^C^*α*^ Chemical Shifts in terms of a single common variance could be miss leading. Instead, we use quantiles computed per each amino acid’s distribution. Specifically we used the 0.05, 0.2, 0.8 and 0.95 quantiles. Thus, we divide the reference densities into a central 60% (between the 0.2 and 0.8 quantiles) a 30% (15% between 0.05 and 0.2 and another 15% between 0.8 and 0.95) and finally the remaining 10% for those value below 0.05 and above 0.95. In Figure 2 we can also see the differences for every residue in structures 1D3Z and 1UBQ represented as rhombuses and dots below each reference density. We use color to help interpret such differences, green if the ^13^C^*α*^ Chemical Shifts difference is found in the central 60%, yellow if it is found in the 30% around the central values and red if it is found in the 10% most extreme values. It is important to note that even when a residue is marked red that does not automatically indicates it is a poorly determined residue, as in fact we expect that 10% of the residues from good quality structures to appear red using the presented method. Instead we can considered them as residues that may deserve further attention [15].

**Figure 2:**
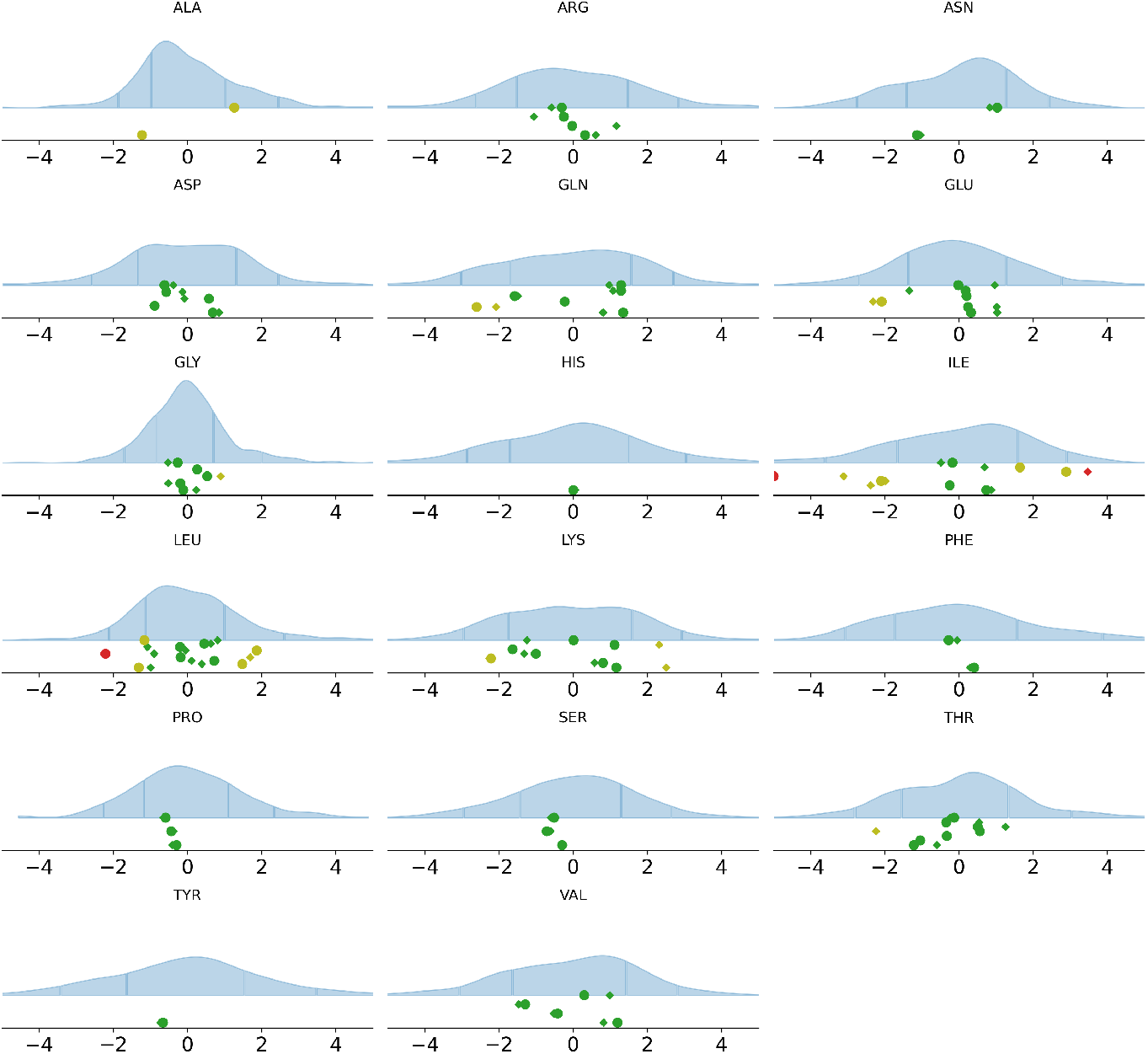
The reference densities in light blue are used as a representation for the expected differences between experimental and corrected ^13^C^*α*^ Chemical Shifts in high quality NMR resolved structures. The dots and rhombuses represent the ^13^C^*α*^ Chemical Shifts differences for the structures under PDB entries 1D3Z and 1UBQ respectively. The markers are displayed in color red when found in the 10% most extreme values, yellow if they are placed in the 30% around the central values and green for the central 60%.

As we can see the differences in general are small between the two target structures, with a few exceptions such as Isoleucine 30 and Leucine 56. Figure 3 uses the same color-schema from Figure 2 in the context of a 3D structure. The accompanying code at the repository (see Abstract) can automatically generate a file containing the colored structures as in Figure 3 and can be loaded with PyMOL or VMD [29].

**Figure 3:**
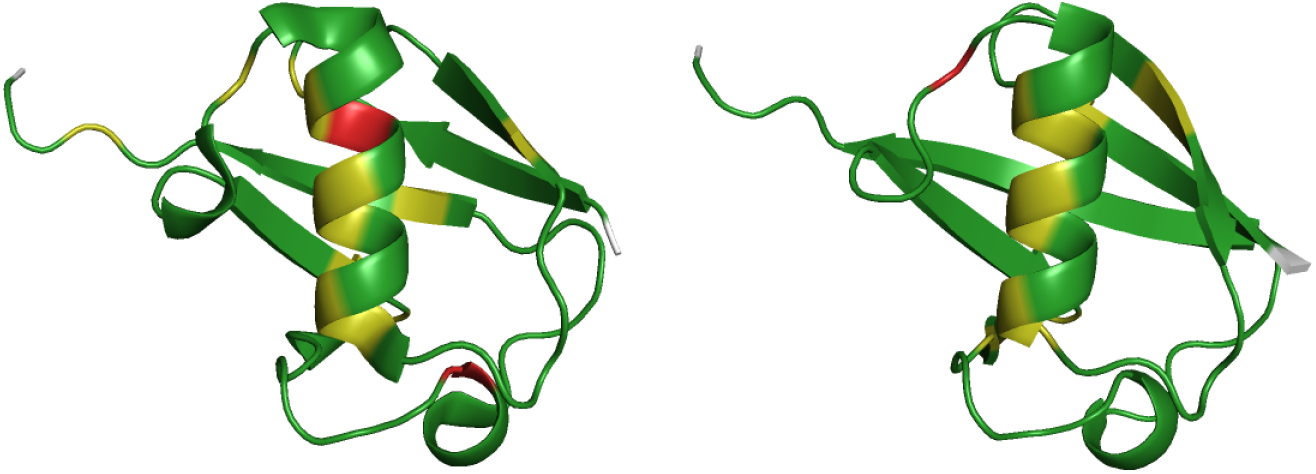
3D structure of 1D3Z (left) and 1UBQ (right). The amino acid residues are colored according to the placement of their ^13^C^*α*^ Chemical Shift difference respective to the reference densities in Figure 2.

### LOO predictive distribution

As previously mentioned the LOO predictive distribution is a general way to assess Bayesian statistical models and is not related to proteins structures in any direct way [11, 17, 18, 25]. For a calibrated model, i.e. a model which predictive distribution is in good agreement with the observed data, the distribution of the LOO probability integral transform (LOO-PIT) is uniform. As this only holds asymptotically, a way to empirically assess calibration for a finite sample is to compare the density of LOO-PIT against the density of uniform samples of the same size as the data used to compute LOO-PIT. Such comparison is done in Figure 4 for structures 1D3Z and 1UBQ. We can see that both models seem to be overall well calibrated but both models have a slightly positive slope, with less predicted lower values than expected and more higher values than expected. In other words the predictive distributions are slightly biased compared to the training data, specially for 1D3Z which it is also showing a small excess of values around the middle.

**Figure 4:**
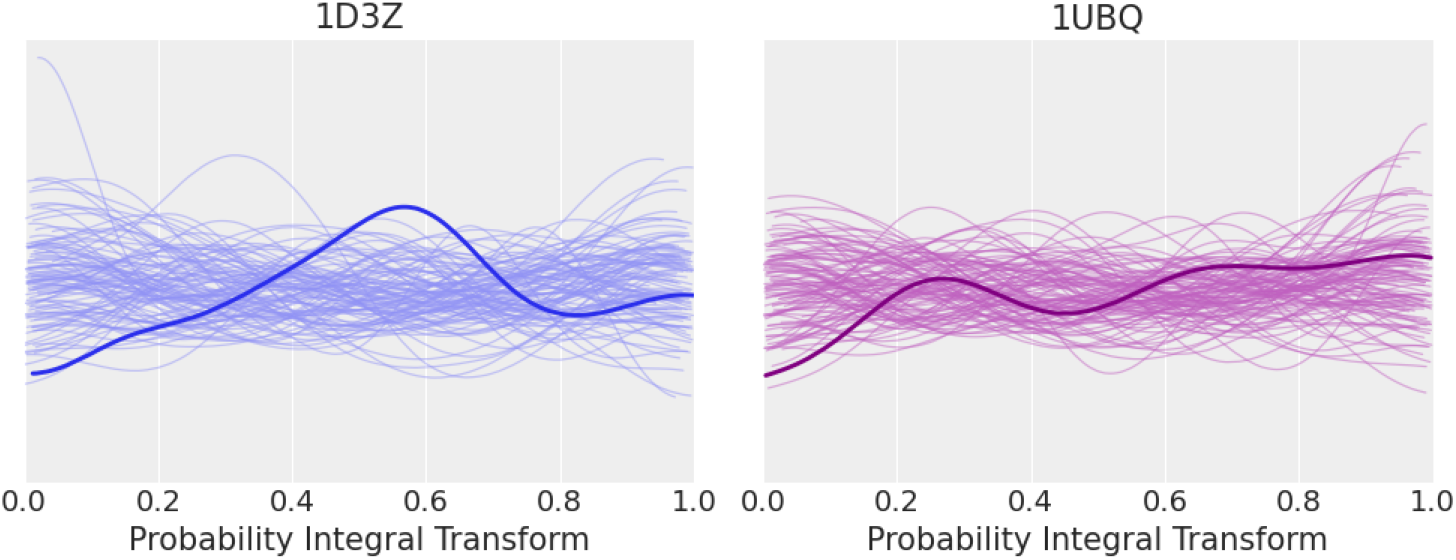
LOO-PIT for 1D3Z and 1UBQ. The thick line corresponds to the observed LOO-PIT density and the thin lines represent simulations from the standard uniform distribution in the [0, 1] interval for a data set of the same size as the one used to computed LOO-PIT. From comparison with these simulations we can define what constitutes a large or small deviation from uniformity. Both 1D3Z and 1UBQ are within the expected margins.

### Analysis of the 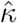 parameter

The parameter 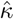 of the Pareto distribution used to stabilize the importance weights during the LOO computation can help spot influential observations, i.e. observations that have a large effect on predictions if removed during the cross validation approximation. The higher this value, the most influential the observation is, with values above 0.7 being of particular interest (see subsection LOO in Methods and Software section). Figure 5 shows that for 1D3Z the residues with 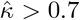 are Asparagine 25, Valine 26, Isoleucine 30 and Isoleucine 61. While for 1UBQ these are Isoleucine 13, Valine 26, Isoleucine 30 and Arginine 72.

**Figure 5:**
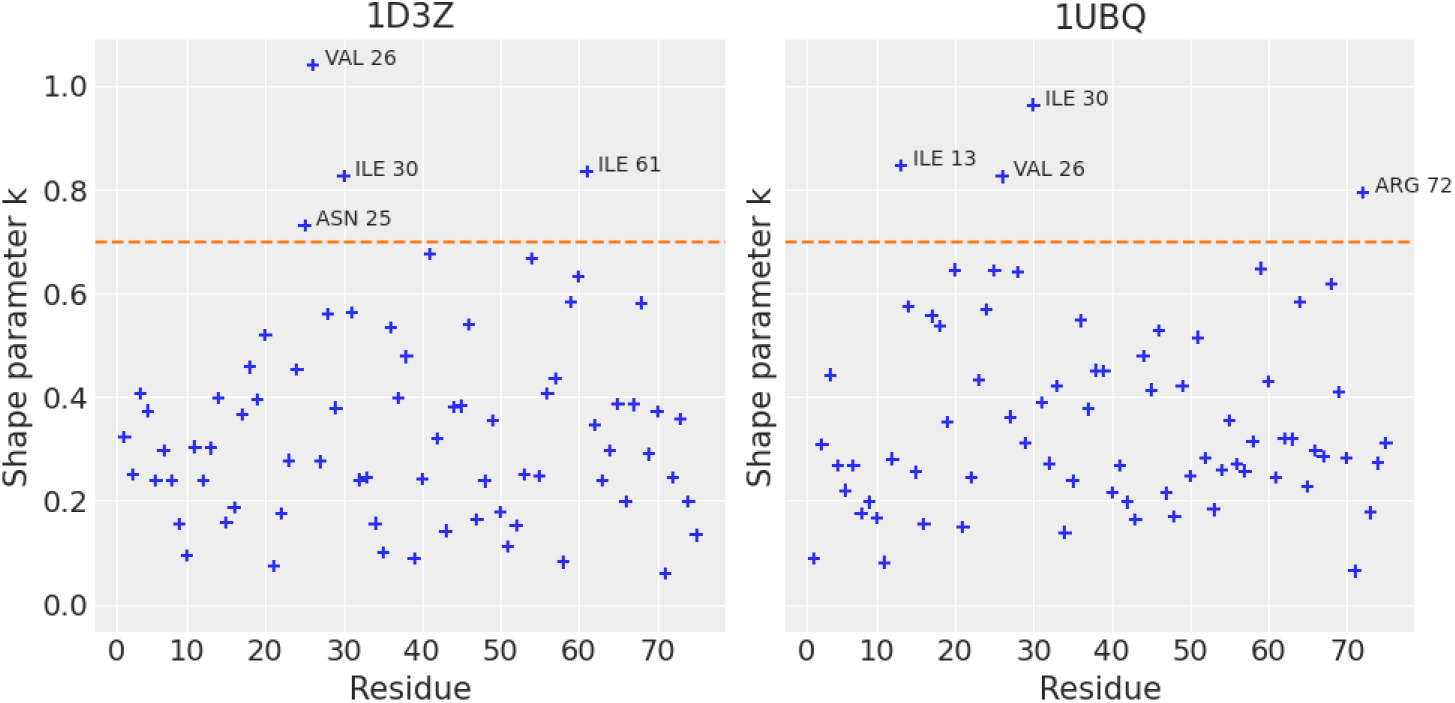
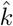 values for every residue in 1D3Z (left) and 1UBQ (right). The dashed orange line indicates the value of 0.7. Residues above this value are labeled as they can be considered as highly influential.

### Expected Log Predictive Density

Figure 6 shows the differences of ELPD between structures 1D3Z and 1UBQ. Globally, these structures seem to be on par, except for residue Isoleucine 30 that is showing a better agreement for 1UBQ.

**Figure 6:**
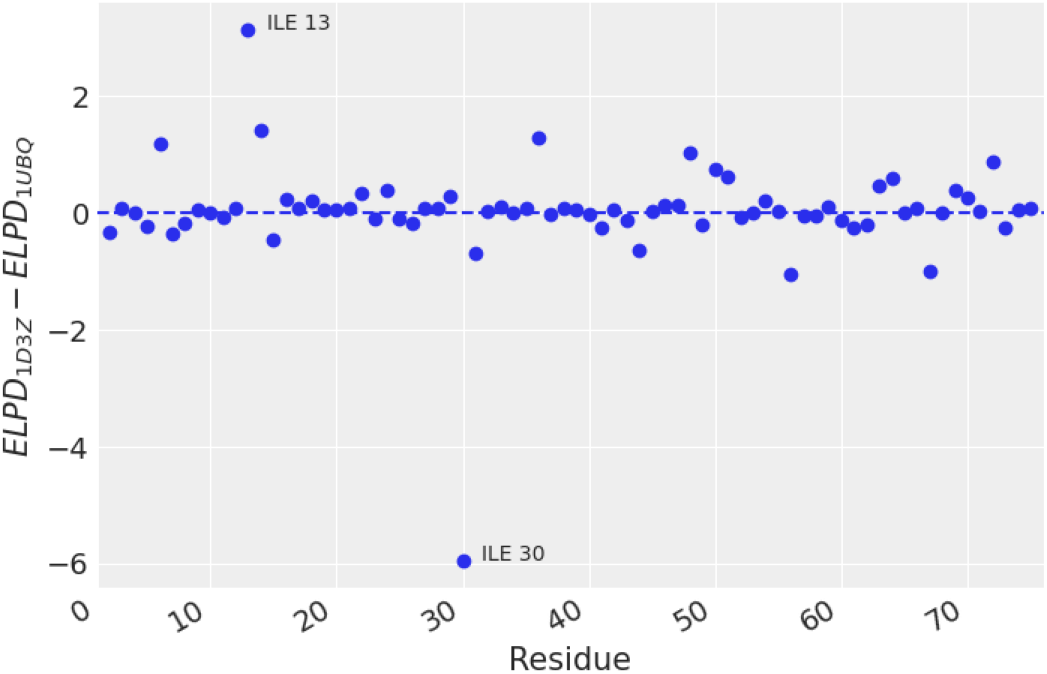
Difference for the Expected Log Predictive Density between structures 1D3Z and 1UBQ. Positive values indicate, that a particular residue is better resolved for structure 1D3Z, and in turn negative values indicate that structure 1UBQ better resolves them.

From the observation of these results, it is worth noting that Isoleucine 30 has a 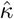 value ≥ 0.7 in both 1D3Z and 1UBQ (see Figure 5). It was also located in the extreme quantiles on Figures 2 and 3 for the 1D3Z target structure. Also, it showed the largest ELPD difference between the two structures analysed in this work (see Figure 6). Another flagged residue for both structures, this time only by the 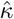 value analysis, is Valine 26. Although, this residue was in the green region of expected differences for both structures. These results perform a demonstration of how the described analyses can be used for structure validation. This implies the use of the presented tools for pointing at residues with poor agreement between experimental and theoretical ^13^C^*α*^ Chemical Shifts according to the Bayesian hierarchical linear model.

## Conclusions

We have presented a collection of tools and visualizations for NMR protein structure assessment. All these tools are based on Bayesian statistical models and established validation methods from that field. We consider such visualizations as useful additions to the current toolbox for protein structure validation. We notice that we are using these Bayesian model comparison tools differently from standard Bayesian model comparison routines. That is to say, the statistical models are kept fixed and the 3D structural models vary. Thus, when observing a potential problem we are directly attributing it to the structure’s quality, as we consider that the hierarchical linear model and the method to compute ^13^C^*α*^ Chemical Shifts are in general good enough for our purposes.

As different observables reveal different aspects of protein structures, we encourage researchers to perform similar analyses to the ones presented here using other observable than ^13^C^*α*^ Chemical Shifts. Finally, we want to emphasize that the tools and visualization presented here are not intended to provide categorical answers about the quality of proteins but instead help experts explore the quality of structures and, when possible, guide them to make improvements of such models.

## Acknowledgments

We are honored to dedicate this manuscript to the memory of Harold A. Scheraga, Professor of Chemistry, Emeritus at Cornell University. He achieved leadership in the world of science, and high respect among colleagues, as a result of his colossal experience in Experimental and Theoretical Chemistry, Physics and Mathematics, research in Protein Chemistry and, in particular, due to his tireless effort in the search of possible solutions to the Protein Folding problem. Harold Scheraga passed away at the age of 98 in Ithaca, NY, on August 1^*st*^, 2020.

Funding was provided by National Agency of Scientific and Technological promotion —ANPCyT, Grant PICT-0767, PICT-02212. And National Scientific and Technical Research Council —CONICET, Grant PIP-0087.)

